# Human-specific GAPDH RT-qPCR is an accurate and sensitive method of xenograft metastasis quantification

**DOI:** 10.1101/2020.07.16.207175

**Authors:** Margaret L Dahn, Cheryl A Dean, Diana B Jo, Krysta M Coyle, Paola Marcato

## Abstract

Metastasis is the primary cause of cancer-related mortality. Having experimental models that accurately reflect changes in the metastatic burden is imperative for developing improved treatments and a better understanding of the disease. The murine xenograft tumor model mimics the human scenario and provides a platform for *in vivo* and *ex vivo* metastasis quantification analyses. Histological analysis of hematoxylin and eosin (H&E) stained thin sections has been the gold standard for quantifying metastasis *ex vivo* but gaining favor for its ease and accuracy is reverse transcription-qualitative polymerase chain reaction (RT-qPCR). Herein we directly compare histological and RT-qPCR-based methods for quantifying lung metastasis in a murine xenograft tumor model. Furthermore, we have introduced a variation of the RT-qPCR method; human-specific glyceraldehyde 3-phosphate dehydrogenase (GAPDH) RT-qPCR, which allows quantification of metastasis in xenograft models, without the requirement of overexpression of exogenous genes. Human-specific GAPDH RT-qPCR detected increased lung metastasis resulting from aldehyde dehydrogenase 1A3 (ALDH1A3) expression in MDA-MB-231 breast cancer cells orthotopically implanted in NOD/SCID mice. Further, in the xenograft tumor model, human-specific GAPDH RT-qPCR was more sensitive and cost-effective than quantification of lung metastasis by histological analysis of H&E stained fixed thin sections. The two assays were highly correlative in terms of determining relative metastatic burden, suggesting that the human-specific GAPDH RT-qPCR method could be used as a standard method for quantification of disseminated human cells in murine xenograft models.

## Introduction

Metastasis is the predominant cause of cancer-related mortality, responsible for approximately 90% of all cancer deaths (1). It is a multistep process wherein cancer cells at the site of origin spread to secondary tissues and organs; the lungs are a common site of metastasis in a wide range of cancers (2). Understanding metastasis and the development of therapeutic agents that prevent metastasis requires *in vivo* metastatic experimental models that reflect the human scenario. Favoured among researchers are murine tumor models of metastasis where cancer cells are orthotopically implanted, allowing for tumor-specific preferences for dissemination to distant sites such as lung, brain, liver, and bone (3). Rapid, cost-effective, and accurate measurement of lung metastasis is important to many pre-clinical projects.

There are numerous quantitative tools to measure metastasis in animal models, some of which are designed to mimic the clinical setting and allow for metastasis quantification in a live animal. These methods include magnetic resonance imaging (MRI) and bioluminescence imaging (BLI) of live animals (4–6). In the clinic, MRI is used to detect, locate, characterize, and stage cancer and assess response to treatment. MRI offers sensitivity and specificity, due to its intrinsic spatial/temporal/contrast resolutions and adequate detectability for tiny amount of substances, which make it ideal for research utilizing *in vivo* models (6). BLI measures photon emission from cancer cells that are engineered to express the luciferase protein and detects the presence of these cells in organs and tissues. The luciferase reaction and production of luminescent product requires live tumor cells and the luciferin substrate that relies on adequate distribution (7). In addition to live animal imaging, BLI can be used *ex vivo* to quantify lung metastasis in dissected lung tissue. Both MRI and BLI require specific costly equipment and facilities, which is prohibitive for some researchers. Hence there is a need for methods that use equipment and facilities common in molecular biology laboratories.

A number of *ex vivo* methods have been developed for metastasis quantification amenable to most molecular biology laboratories. Flow cytometry analysis can be used for lung metastasis quantification. For example, cancer cells engineered to express green fluorescent protein (GFP) can be detected by flow cytometry and relative percentages of GFP+ cells determined in harvested tissues (8). However, this method is likely error prone as red blood cell-free single-cell suspensions need to be generated and the cancer cells need to be tagged with or express a fluorochrome which could alter cell behaviour. Alternatively, clonogenic based-assays can be used to quantify lung metastasis (9–12). In this technique, harvested tissues are dissociated into a single-cell suspension, plated as single cells and cultured for at least 10-14 days in a drug selection media (e.g. 4T1 cells are resistant to 6-thioguanine (13)), followed by subsequent staining and enumeration of resulting colonies. This assay requires a method of selection of cancer cells over non-cancer cells, which limits the models that can be used and is potentially error prone (since it relies on the assumption that the colonies that form from the processed samples accurately represent the metastasis that was present in the tissues at the time of harvest).

Stereological analysis estimates metastasis volumes in mouse lungs by hematoxylin and eosin (H&E) staining of thin sections of cryostat embedded lungs (14). A variation of the stereological analysis is to perform H&E staining of thin sections of fixed, paraffin embedded tissues and perform microscopic quantification of metastatic surface areas. This closely mimics the clinical pathological practice and it is commonly used and is therefore often seen as the gold standard method (15–20); however, achieving quantitative values of metastasis by histological examination is labor intensive and requires quantification of thin sections throughout the specimen (21).

An alternative *ex-vivo* method gaining favour is based on the amplification of tumor-specific transcripts using reverse transcription-quantitative polymerase chain reaction (RT-qPCR). This method gives an assessment of the overall metastatic burden of the total tissue, independent of uneven distribution of metastasis in the tissue. RT-qPCR has been shown to be a sensitive method, detecting 10 cancer cells per 0.5-2 μg of lung tissue (22). However, this was based on detection of overexpressed transcript found in only some cell lines (i.e. the human epidermal growth factor receptor 2, HER2, transcript in HER2 overexpressing cell line JIMT1). Similarly, others have been able to quantify lung metastasis by RT-qPCR by ectopic overexpression of a foreign gene (i.e. luciferase gene) (23). Luciferase gene expression quantification by RT-qPCR in harvested lungs was shown to be correlate with metastasis quantification by BLI and is 10 times more sensitive, suggesting that RT-qPCR could be used as an endpoint readout when used in combination with BLI.

The wide spread use of RT-qPCR as a technique for evaluating murine metastasis is thus limited by the requirement of a highly expressed gene that is unique to the cancer cells and the lack of comparison between metastasis quantification by RT-qPCR and a well-accepted method (e.g. histological analysis of H&E stained thin sections). Herein we describe our method of quantifying expression of abundant human and mouse glyceraldehyde 3-phosphate dehydrogenase (GAPDH) transcript by RT-qPCR utilizing validated human-specific primers. We demonstrate that human-specific GAPDH RT-qPCR is highly sensitive, discriminating between transcripts of human and mouse origin with high specificity. The method reliably detects 100 human cells in a mouse lung lobe (~70mg tissue), equating to a detection limit of ~1 human cell/0.7mg of mouse lung tissue). Human-specific GAPDH RT-qPCR was successfully utilized to detect increased metastatic burden imparted by aldehyde dehydrogenase 1A3 (ALDH1A3) expression in MDA-MB-231 cells. A direct comparison of lung metastasis quantified using human-specific GAPDH RT-qPCR versus histological quantification of H&E stained thin sections and found the two methods highly correlative, with the former being more sensitive.

## RESULTS

Herein we compare two methods for quantification of lung metastasis in a murine xenograft tumor model; the gold-standard histological based method of quantifying percent metastasis of sectioned and stained fixed lung tissue versus an RT-qPCR-based method of quantifying levels of human cancer cell-specific transcripts in RNA extracted from murine lung tissue. We have introduced a variation of the RT-qPCR method that will allow accurate and sensitive quantification of metastasis without the requirement of overexpression of exogeneous genes in the context of xenograft models (i.e. human-specific GAPDH RT-qPCR). In Figure 1, we outline the workflow and how we will directly compare these two methods using the same experimental lung specimens.

**Figure 1.**
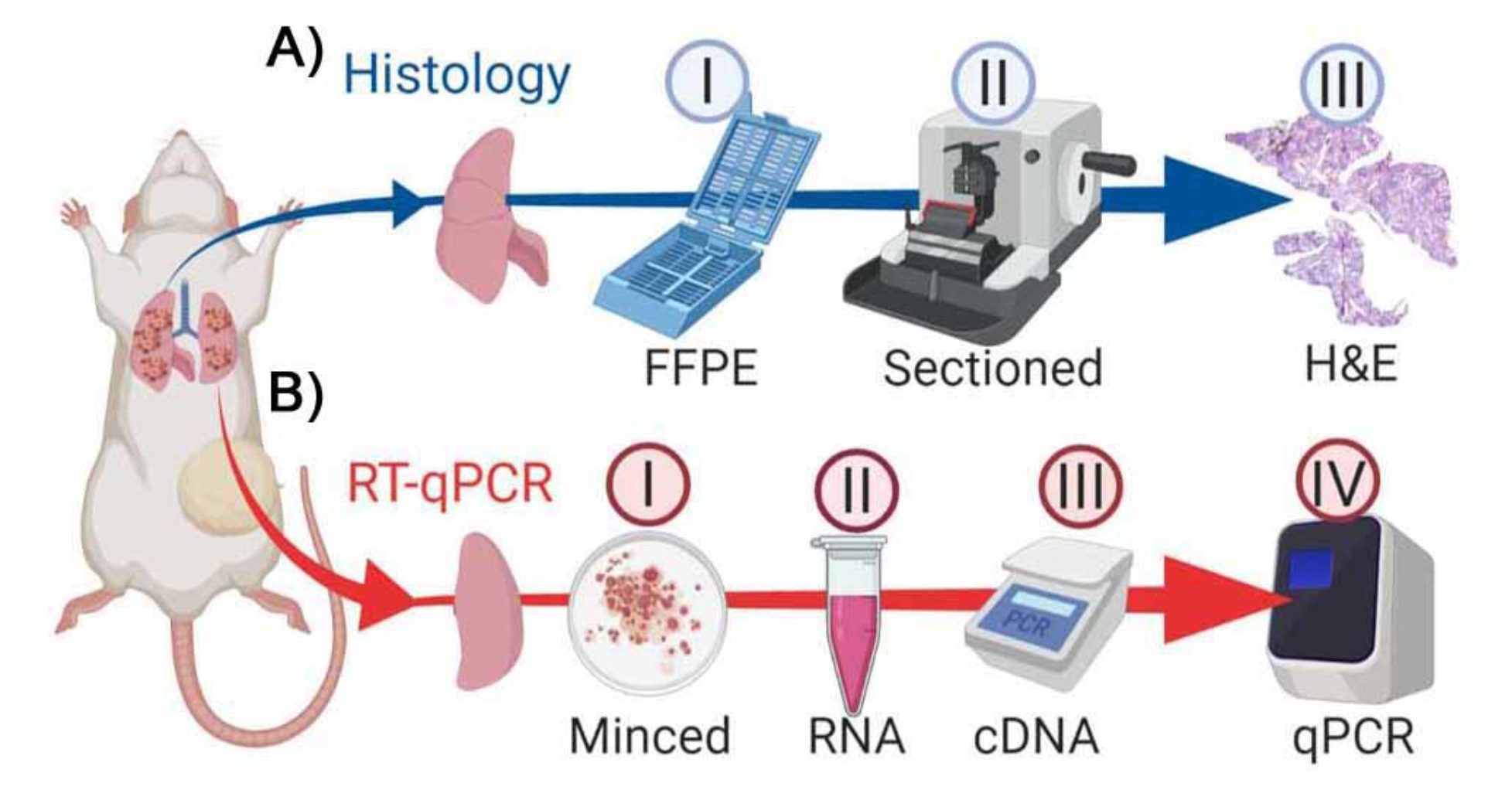
Schematic representation of workflow comparing quantification of lung metastasis by histological analysis of H&E stained fixed thin section versus human-specific GAPDH RT-qPCR. **A)** Lung processed for histology-based quantification. I) Formalin-fixed paraffin-embedded (FFPE): majority of multi-lobed lung kept in histology cassette for 24 hours in formalin then stored in 70% ethanol followed by dehydration/clearing/embedding. II) Sectioning by microtome cutting. III) H&E staining includes deparaffinization/rehydration/ staining followed by dehydration and cover slipping. Images were captured by at 2.5x magnification and metastasis quantified in Image J. **B)** Lung processed for RT-qPCR-based quantification. I) Single-lobed left lung minced/homogenized and 70mg added to II) 1mL of Trizol for subsequent RNA extraction. III) cDNA synthesized and IV) qPCR performed with GAPDH human-specific primers and non-specific mouse GAPDH primers as a reference gene.

### Histological quantification of lung metastasis

To compare the two methods of metastasis quantification, we established a tumor xenograft model with previous demonstrated differing levels of lung metastasis to better test the sensitivity of the methods. We utilized MDA-MB-231 cells, with or without ALDH1A3 overexpression (20). Notably, by using histological quantification methods we previously determined that when these cells are orthotopically implanted in the mammary fat pads of NOD/SCID mice, lung metastasis occurs in approximately 50 days and that ALDH1A3 overexpression increases the metastatic burden (20). Here, we terminated the experiment on day 49, noting a significant increase in tumor volume upon ALDH1A3 overexpression (OE, Figure 2A). We determined the metastatic burden resulting in each mouse lung using the gold-standard method of quantification (histological analysis of H&E stained lung thin sections spaced throughout the fixed lung paired with Image J analysis of captured images, Figure 2B, Supplemental Figure 1). This established a reference for later comparison with the RT-qPCR method and confirmed that the MDA-MB-231 cells had metastasized to the lungs in these specimens and that ALDH1A3-OE increased metastasis (Figure 2C). Using the histological analysis method, we detected lung metastasis in 5/15 of the control and 11/11 ALDH1A3-OE lung specimens (16/26 lungs total). In the 16 positive metastatic tissue specimens, metastasis ranged from 0.07% to 92.14%. Therefore, histological quantification of the lungs discriminates differences in metastatic burden between different experimental conditions and the limit of detection of metastasis was 0.07% of total lung tissue (averaged over 4 lung lobes and multiple thin sections/specimen).

**Figure 2.**
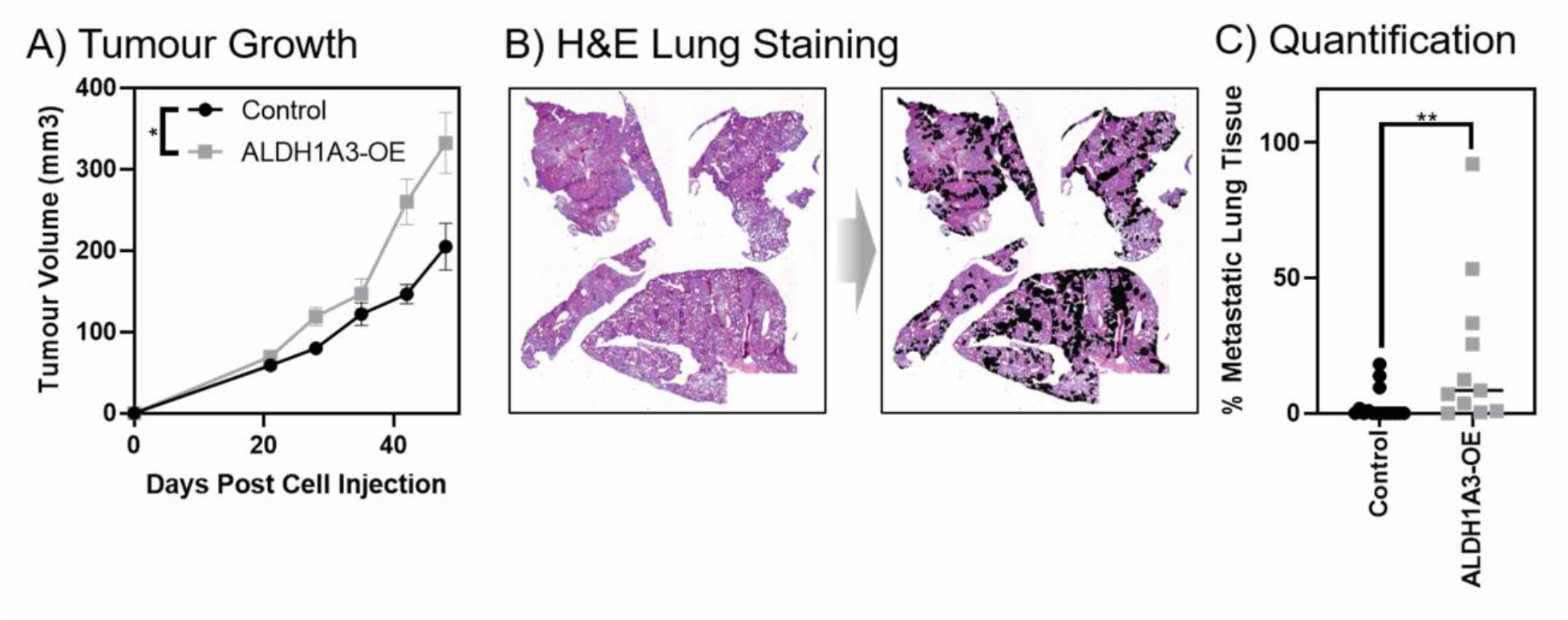
Histological quantification by H&E staining of fixed thin sections demonstrates that ALDH1A3 expression increases metastasis to the lungs of orthotopically established MDA-MB-231 tumors. **A)** ALDH1A3 overexpression increases MDA-MB-231 tumour volume and (Control n=15, ALDH1A3 overexpression n=11, SEM error bars). **B)** Light microscopy image of H&E stained lung section; example from ALDH1A3 overexpression tumour bearing mouse with metastatic nodes outlined. **C)** Histology quantified lung metastasis (median, SEM error bars); p<0.05*.

### Identification of human-specific GAPDH primers that discriminate between mouse tissue and human cells

High sensitivity in RT-qPCR based methods of lung metastasis quantification is more likely achievable if detecting abundant cancer cell-unique transcripts. GAPDH is among the most abundantly expressed transcripts; hence, if human-specific primers to GAPDH are successfully designed that would discriminate between human and mouse transcripts in RT-qPCR, it could be a highly sensitive method for detection of disseminated human cancer cells in murine xenograft models. Further, it would preclude the requirement of introduction of a foreign gene and be amenable to all xenograft models, eliminating the need for a cell line-specific overexpressed gene (e.g. HER2).

Using NCBI Primer-BLAST, we designed primers to unique regions of human GAPDH transcript that would generate amplicons of 86bp for the human GAPDH transcript (Supplemental Table 1). The primers have a theoretical annealing temperature of 60°C. We performed an *in silico* analysis of the specificity of the human GAPDH primers against the human and mouse transcriptome. We set very wide parameters for the theoretical RT-qPCR products that could be generated with a wide primer annealing temperature window (57 - 63°C) and large amplicon size range (70 - 1 000 bp) to capture any potential amplification, no matter how unlikely. As expected, when we ran the human-specific primers against human reference transcriptome (NCBI RefSeq Database) there is 100% fidelity for GAPDH TV1-4 for both primers. There are two mismatches for the forward primer with TV7. When we ran the human-specific primers against the mouse reference transcriptome it revealed one potential unlikely off-target with mouse myosin phosphatase rho Interacting protein (MPRIP), with 5 mismatches for the forward primer and 4 mismatches with the reverse primer.

We used the human GAPDH primers in RT-qPCR experiments against RNA isolated from cultured MDA-MB-231 cells and naïve NOD/SCID mouse lung (harvested from a mouse that had no exposure to human cells). We similarly used mouse GAPDH primers against the two different templates. Serial dilutions of the cDNA templates in the qPCR reactions revealed that our human GAPDH primers were highly specific to the template of human origin (MDA-MB-231 cells) (Figure 3A). There was no product when the human GAPDH primers were used in qPCR reactions with mouse lung cDNA template. In contrast, the mouse GAPDH designed primers amplified a product to cDNA templates of both human (MDA-MB-231 cells) and mouse origin (mouse lung tissue, Figure 3A). The data with the mouse primers reinforces the notion that all primers made to a certain species need to be validated in RT-qPCR to confirm the primer specificity for the template of said species. The melting curves confirmed that only one product was consistently made in the qPCR reactions, even in reactions with highly diluted template (Figure 3B). Using the amplification data from the serially diluted cDNA qPCRs (Figure 3A), we generated standard curves (Figure 3C). This revealed that the primers were highly efficient over a wide range of cDNA template concentrations (Figure 3C). Finally, we tested the primers in side-by-side reactions with three different samples of concentrated MDA-MB-231 and mouse lung (Figure 3D). This confirmed that the human-specific GAPDH primers generate amplicons from templates of human origin and not mouse, while the mouse primers are non-specific in terms of species (Figure 3D). Together, this data validated that the human GAPDH primers are highly human-specific and efficient in terms of generating a product by RT-qPCR (Figure 3). Importantly, our metastasis quantification assay requires that only the human GAPDH primers be highly specific to human cDNA template; therefore, we proceeded to the next validation step.

**Figure 3.**
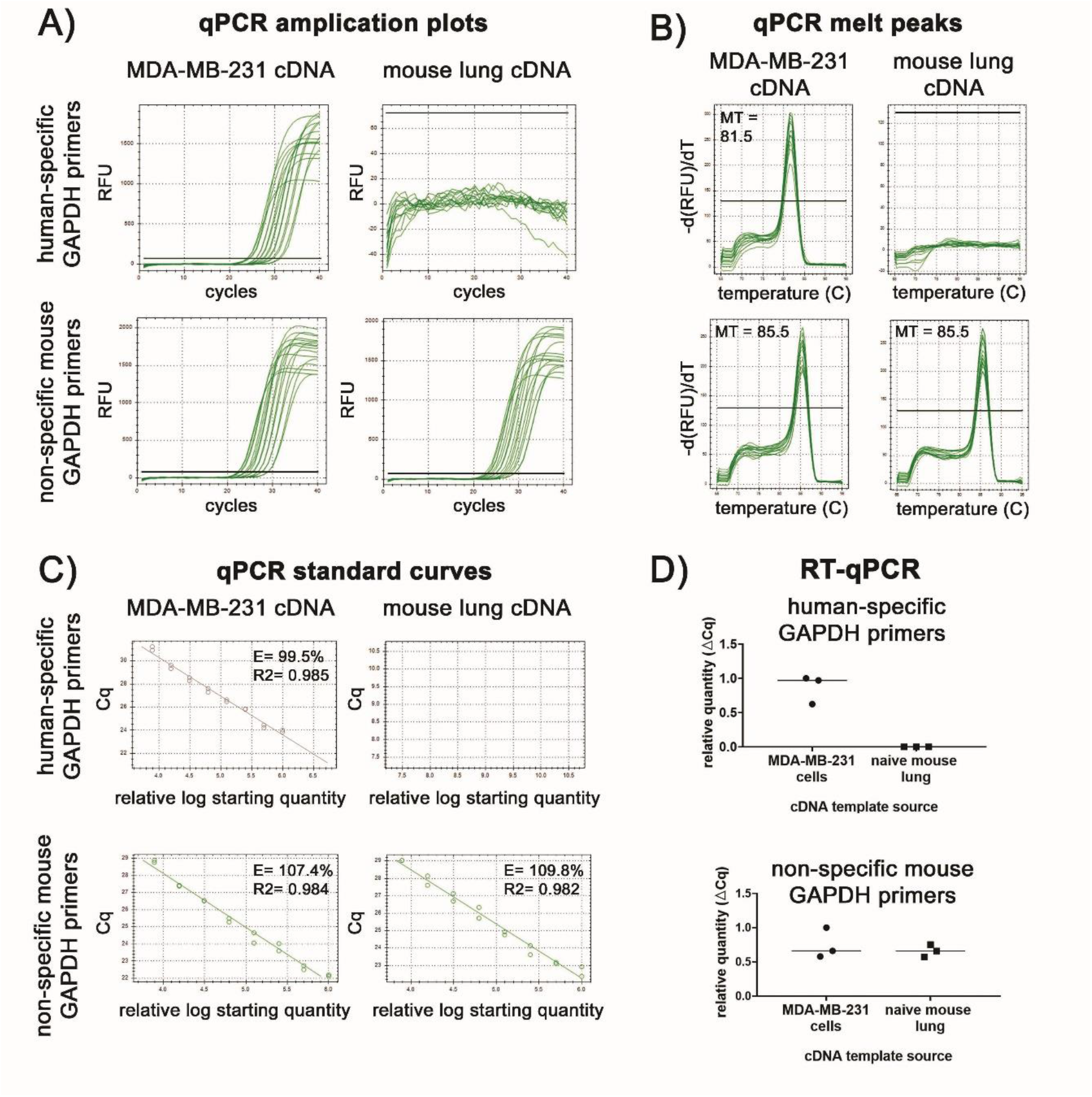
Validation that the human GAPDH primers are efficient human-specific primers. The qPCR amplification curves **(A)** and melt curves **(B)** generated by the human and mouse GAPDH primers using cDNA template from RNA extracted MDA-MB-231 cells and naïve NOD/SCID mouse lung. MT = melting temperature in Celsius at the peak. **C)** Standard curves were generated using data from **(A)**. E = efficiency and R2 = the square of the correlation (the coefficient of determination). **D)** Three different concentrated MDA-MB-231 and naïve NOD/SCID mouse lung RNA samples were run RT-qPCRs with the human and mouse GAPDH primers.

### Human-specific GAPDH primers identify as few as 100 human cells in a mouse lung lobe, with a linear range between 100 – 1 000 000 human cells in the context of mouse lung

Having identified human GAPDH primers with high specificity for transcripts of human origin, we next determined the detection and specificity limit of the human GAPDH primers for human GAPDH transcripts in the context abundant mouse transcripts. For this analysis we added known amounts of serially diluted MDA-MB-231 cells (0, 10, 100, 1 000, 10 000, 100 000, 1 000 000 cells) into tubes containing a naïve NOD/SCID lung lobe. We then purified the total RNA and performed RT-qPCR on the samples using the human-specific GAPDH primers and non-specific mouse GAPDH primers as a reference gene. The resulting CT values demonstrate the high specificity of the human-specific GAPDH primers; in the context of abundant mouse RNA, the human GAPDH transcript was detectable when as few as 100 MDA-MB-231 cells were spiked into the mouse lung lobe sample (Figure 4A). The standard curve revealed that quantification of human GAPDH transcript in the context of abundant mouse lung RNA was linear between the range of 100 to 1 000 000 MDA-MB-231 cells (Figure 4B).

**Figure 4.**
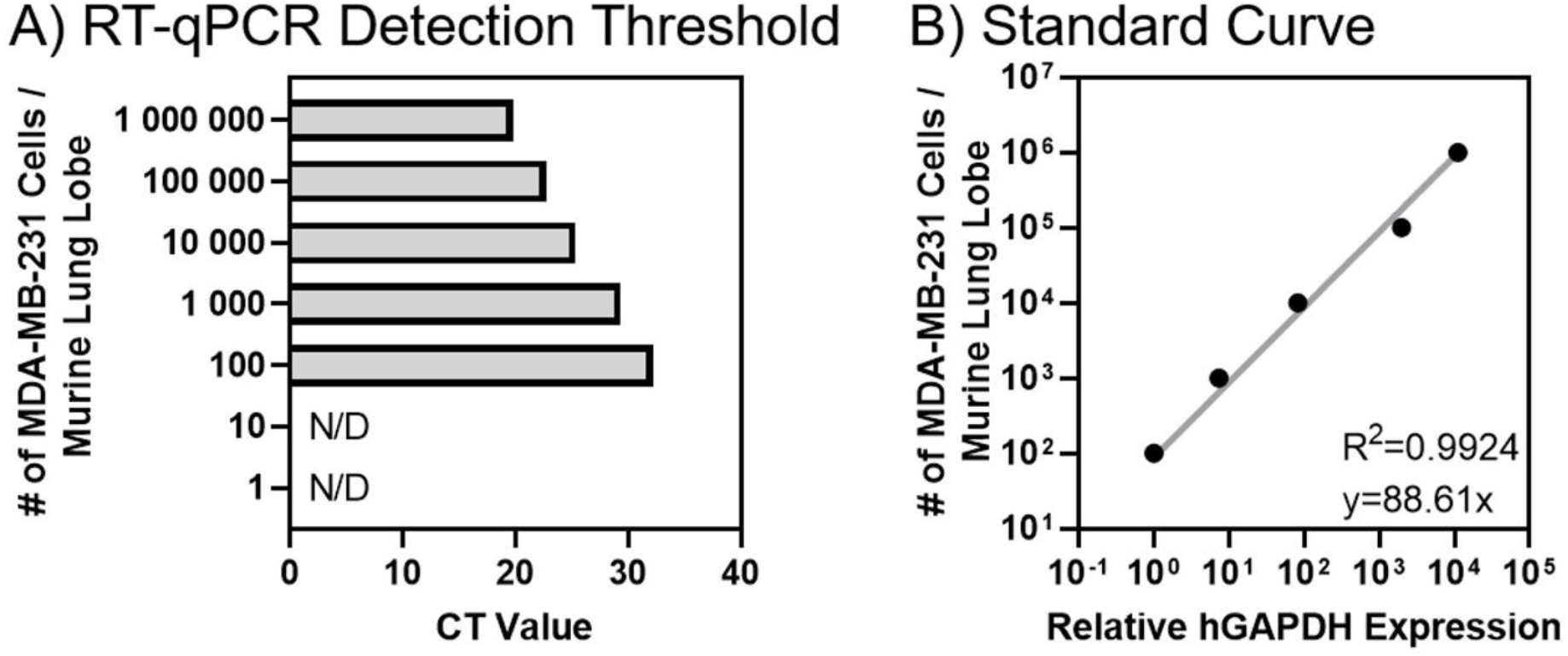
The detection limit of human-specific GAPDH RT-qPCR is 100 human cells per mouse lung lobe and the standard curve is linear range between 100 – 1 000 000 human cells per mouse lung lobe. **A)** Increasing numbers of MDA-MB-231 cells were added to naïve NOD/SCID lung lobe samples and total RNA extracted and the detection limit of human-specific GAPDH primers determined by RT-qPCR. N/D = not detected. **B**) Standard curve of the number of MDA-MB-231 cells added in naïve NOD/SCID lung lobe plotted against the relative amount of human GAPDH transcript detected by RT-qPCR. R^2^ = the square of the correlation (the coefficient of determination). The human GAPDH (human-specific primers) per each sample was made relative the detected total GAPDH (non-specific mouse primers).

### Human-specific GAPDH RT-qPCR is highly sensitive and an accurate method of xenograft lung metastasis in comparison to histological quantification

Having established the dynamic range and specificity of the human-specific GAPDH RT-qPCR, we next quantified the lung metastasis in the left lobe of the harvested lung from the MDA-MB-231 xenograft experiment (with or without ALDH1A3 overexpression, Figure 2A). Using the standard curve (Figure 4B), this revealed the number of MDA-MB-231 cells present in each lung lobe sample. All the experimental lung samples fell within the standard curve (Figure 5A) and were therefore usable samples for metastasis quantification. As expected, based on the histological analysis of the same lung specimens (Figure 2), human-specific GAPDH RT-qPCR also showed that ALDH1A3 overexpression resulted in increased metastasis to the lungs of the mice (Figure 5B). Using this RT-qPCR method, we found between 71 to 219 750 MDA-MB-231 cells in every lung lobe sample; in comparison to the histological method, no metastasis was detectable in 10 lungs (Figure 6A). This demonstrates that human-specific GAPDH RT-qPCR is more sensitive than histological quantification of lung metastasis in the xenograft tumor model. Finally, we determined that the two assays are highly correlative (Figure 6B), demonstrating that human-specific GAPDH RT-qPCR is an accurate method of lung metastasis when directly compared to the gold-standard histological method. Of note is that the correlation is negatively impacted by samples with less than 1% lung metastasis due to those samples falling near the detection limit of metastasis for the histological quantification method.

**Figure 5.**
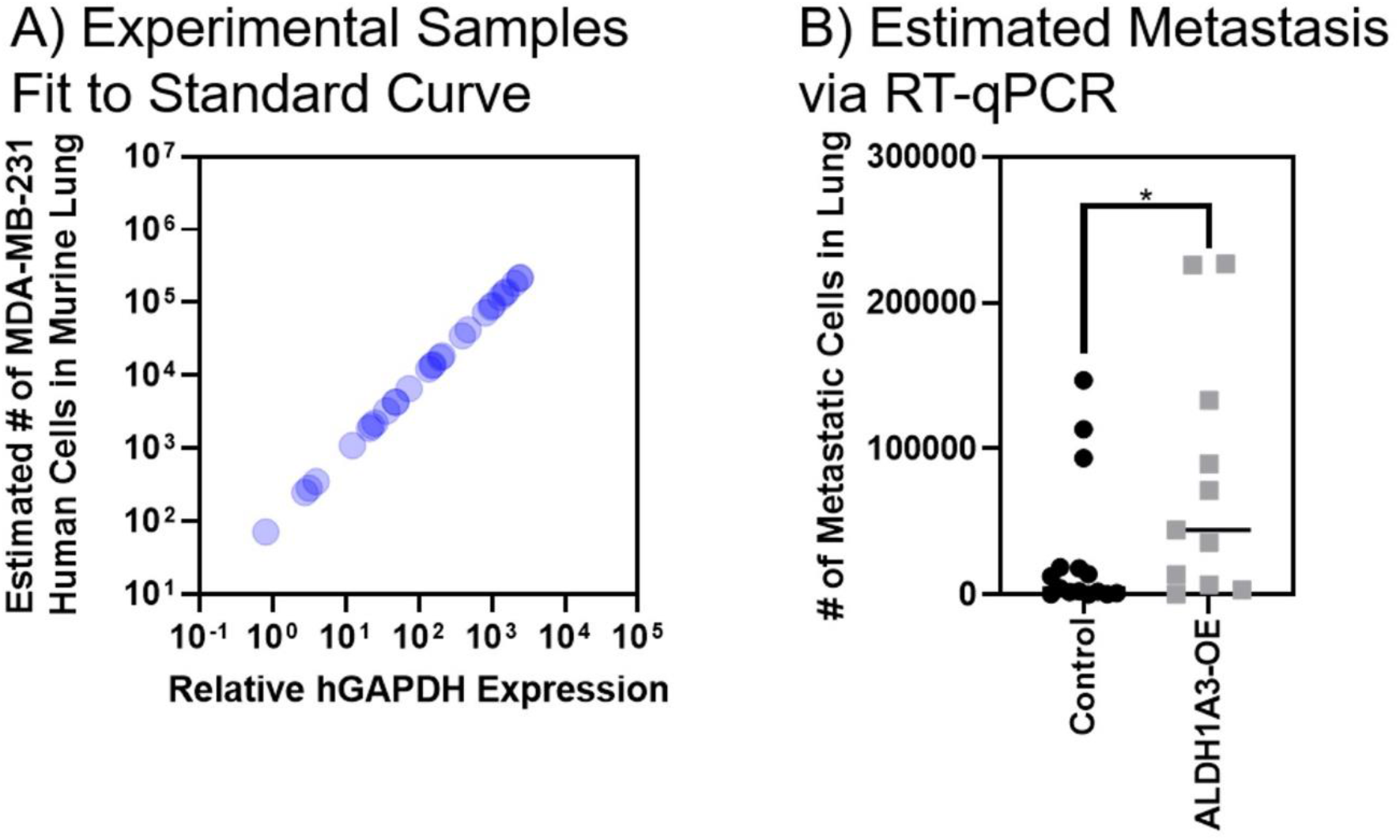
Human-specific GAPDH RT-qPCR quantification of lung metastasis demonstrates that ALDH1A3 expression increases metastasis to the lungs of orthotopically established MDA-MB-231 tumors. **A)** The number of MDA-MB-231 cells per lung lobe is the experimental samples is determined by using the standard curve. **B)** The number of MDA-MB-231 cells/lung lobe in control (n=15) versus ALDH1A3 overexpression (n=11) samples is compared by human-specific GAPDH RT-qPCR median, SEM error bars); p<0.05* and determined using the standard curve of known MDA-MB-231 cells/lung lobe. The human GAPDH per each sample was made relative the detected mouse GAPDH.

**Figure 6.**
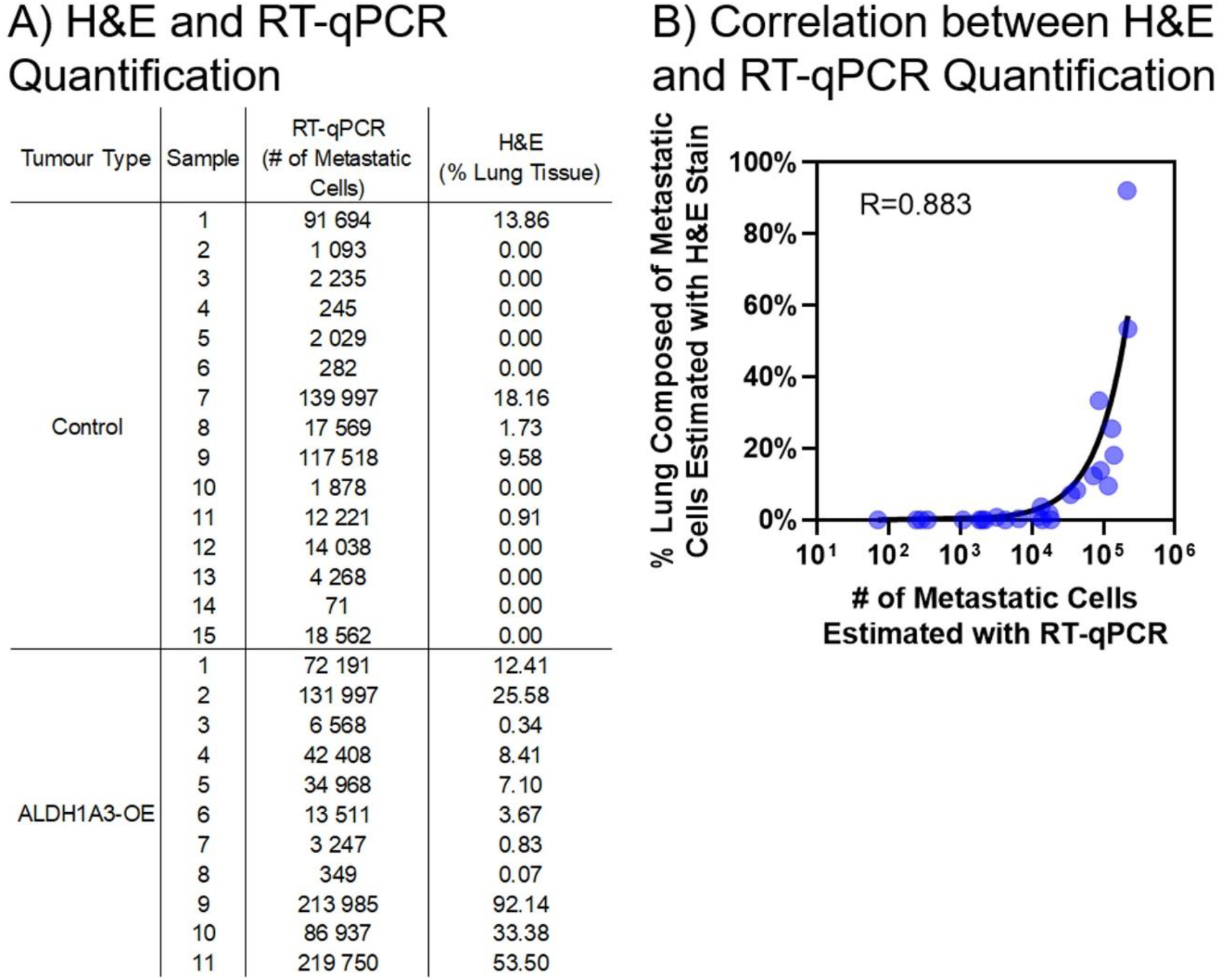
Human-specific GAPDH RT-qPCR correlates with and is more sensitive than histological quantification of lung metastasis by H&E staining of fixed thin sections. **A)** Summary of the lung metastasis quantified in all the experimental samples determined by human-specific GAPDH RT-qPCR (left, # of MDA-MB-231 cells/lung lobe) versus histological quantification by H&E staining of thin sections (right, % of MDA-MB-231 metastasis area averaged over 4 lobes of lung tissue). **B)** The lung metastasis values determined in every experimental samples by human-specific GAPDH RT-qPCR (x-axis) versus histological quantification of H&E stained fixed thin sections (y-axis) are plotted and the Pearson correlation (r) is calculated.

### Human-specific GAPDH RT-qPCR is a relatively cost-effective and rapid method of xenograft lung metastasis

Based on our comparison of the various methods of murine metastasis quantification, human-specific GAPDH RT-qPCR presents several advantages, including its relative low cost and quick method. A time and cost comparison of histological/H&E thin section staining versus human-specific GAPDH RT-qPCR methods show that the former is much more labour-intensive. To quantify lung metastasis by the histological approach, it required approximately 120 hours of technician time to perform lung dissection, tissue embedding and sectioning, H&E staining, image capturing, image stitching and Image J analysis. In contrast, human-specific GAPDH RT-qPCR on the same number of samples required approximately 25 hours of technician time to perform lung dissection, RNA extraction and quantification, cDNA generation and RT-qPCR analysis. This equates to approximately 21% of the time to obtain quantification of lung metastasis by RT-qPCR versus histological analysis, which translates to having the results in less than a week using the RT-qPCR method, and nearly one month using the histological quantification method. Therefore, despite the perhaps increased cost of reagents for RT-qPCR over histological analysis, when the personnel time is considered, RT-qPCR is the more economical and efficient option.

## DISCUSSION

Herein we have described and directly compared the use of human-specific GAPDH RT-qPCR and histological analysis of H&E stained fixed thin sections to quantify lung metastasis in a murine xenograft tumor model. We provide instruction for dissecting and sectioning of mouse lung tissue, utilizing the left lobe for RT-qPCR, leaving the 4 right lobes for alternate analyses if required; our procedure utilized the 4 remaining lobes for histological quantification. Several modifications have been provided for lung dissection, RNA extraction, human-specific primer generation and RT-qPCR that differentiate and improve upon methods reported previously (5, 22).

We acknowledge that other methods, including continuous monitoring of live animals (possible with BLI and MRI) and end-point analyses (e.g. clonogenic assays, flow cytometry and stereological analyses), are also effective for detecting/quantifying metastasis, each having their own strengths and limitations. Stereological and histological analysis by H&E staining closely mimic the clinical pathological practice and are therefore commonly used to quantify lung metastasis, with the latter being the gold standard (24, 25). However, histological examination of the whole organ (i.e. multiple sections from throughout the fixed tissue) is necessary to avoid variability due to sampling error and is largely qualitative unless an imaging software for quantification is also incorporated (21). Therefore, it becomes a labour-intensive procedure. In addition, the smallest metastases that can observed reliably contain ~10 cells in a foci (14).

RT-qPCR has proven to be a sensitive measure to quantitatively detect cancer cell transcripts (exogenously introduced or overexpressed in a specific cancer cell line) within the mouse lung tissue (22, 23). Although RT-qPCR lacks the ability to differentiate between living and dead tumor cells, it provides an accurate and sensitive method for quantification of lung in murine metastasis tumor models. Human-specific primers have been utilized in qPCR to quantify mouse cell contamination in human xenografts (26). Alcoser et al. generated prostaglandin E receptor-2 (PTGER2) human-specific primers to detect contaminating mouse DNA in multiple tumor types (26). Here we performed human-specific GAPDH RT-qPCR on purified RNA instead of qPCR on purified DNA as performed by Alcoser et al. (26). The benefit of human-specific GAPDH RT-qPCR of purified RNA over human-specific qPCR of purified DNA is the much greater template copy number in the former method. This makes RT-qPCR with purified RNA more sensitive over qPCR with purified DNA with respect to detecting a few human cancer cells in abundant mouse tissue (i.e. 1 human cell/0.7mg of mouse tissue).

To our knowledge, no other groups have utilized human-specific primers designed to unique regions of abundant human transcripts to quantify lung metastasis in murine xenograft tumor models. We demonstrate that human-specific GAPDH RT-qPCR is highly sensitive and an accurate method of detecting xenografted cancer cells that have disseminated in mice. Importantly, this method precludes the requirement of the introduction of a foreign gene, or a gene that maybe overexpressed in only some cancer cell lines (e.g. HER2, (22)). This makes human-specific GAPDH RT-qPCR amenable to all xenograft models without consideration for identifying an overexpressed gene or manipulation of the cell line. Perhaps the best indicator of increased sensitivity of the human-specific GAPDH RT-qPCR is the detection of cancer cells in 10/25 experimental samples that were assessed as lacking metastatic cells by H&E staining of fixed thin sections.

In addition to increased sensitivity and specificity, other advantages of this RT-qPCR method include its usability in conjunction with other quantification methods, such as an end point assay when paired with methods such as MRI/BLI quantification of metastasis of live animals. RT-qPCR is also more time/cost effective than many other methods. Furthermore, the method lends itself to quantification of disseminated cancer cells in other tissues besides the lung. For example, it could be used to detect the presence of cancer cells in the lymph node, liver, or brain, and quantify circulating tumor cells.

We propose that RT-qPCR analysis can be performed in lieu of the other described metastasis quantification models. Importantly, the demonstration here that human-specific GAPDH RT-qPCR correlates highly with the gold-standard histological method, is critical if the RT-qPCR method is to be used with confidence over other accepted metastasis quantification methods. Furthermore, our inclusion here of the ALDH1A3 overexpression model was to test the capacity of the RT-qPCR method for discriminating between different conditions with varying metastatic burden. Both the human-specific GAPDH RT-qPCR method and the histological method similarly indicated that ALDH1A3 overexpression in MDA-MB-231 cells increased lung metastasis (p value < 0.05). These analyses will be valuable for other researchers considering RT-qPCR as a primary method of metastasis quantification. RT-qPCR as method of metastasis burden in murine xenograft models may be best suited for research groups with limited access to certain types of equipment and have time and budget constraints.

In conclusion, the data presented suggests that human-specific GAPDH RT-qPCR provides an efficient, sensitive, and cost-effective method of detecting human tumor-derived mRNA within murine tissues. The protocol should be considered as a primary method of metastasis quantification when utilizing murine xenograft models.

## EXPERIMENTAL PROCEDURES

### Cell lines

MDA-MB-231 cells were obtained from the American Type Culture Collection and cultured at 37°C in a 5% CO_2_ incubator. MDA-MB-231 cells were cultured in Dulbecco’s Modified Eagle Medium (DMEM, ThermoFisher, Life Technologies, Cat# 12430062) supplemented with 10% fetal bovine serum (FBS, ThermoFisher, Life Technologies, Cat# 12483020) and 1X antibiotic-antimycotic (ThermoFisher, Life Technologies, Cat# 15240062). Aldehyde dehydrogenase 1A3 (ALDH1A3) overexpressing and control MDA-MB-231 clones were previously generated (20) and maintained in media supplemented with 0.25μg/mL puromycin (Sigma-Aldrich, cat# P8833).

### Murine xenograft metastasis model

A xenograft model was utilized as previously described (20). Briefly, two groups of NOD/SCID female mice were orthotopically injected in the mammary fat pad #4 with 2×10^6^ cells of either ALDH1A3 overexpressing or vector control MDA-MB-231 cells (admixed with high concentration Matrigel, BD, Cat # 354262). Mice were euthanized 49 days post cancer cell implantation and the lungs were harvested. All animal experiments comply with the standards set by the Canadian Council on Animal Care and were performed according to a protocol approved by Dalhousie University’s Committee on Laboratory Animals (protocol 19-013).

### Mouse dissection to harvest lungs

Methods for mouse dissection have been previously published in detail (22); however, we have made some modifications to the protocol, incorporating details described by Sleigh et al. (2016) (27) for removing the pelts. Mice were doused with 70% ethanol to allow for ease of cutting and to sterilize mouse. With sterile dissecting scissors, a small incision was made in the dorsal skin at the level of the hips and continued horizontally around the whole mouse. The pelt was pulled up and over the head of the mouse to expose chest and rib cage. The peritoneum was cut just under and across whole rib cage, and then vertically up with middle of the rib cage. Scissors were then used to cut open the rib cage to expose lungs. The lungs were carefully removed using scissors and forceps. Thymus, heart, efferent and afferent blood vessels, trachea, and connective tissue were removed from the lung tissue. The left lung lobe was separated from the rest of the lung tissue (Supplemental Figure 1). The lungs were rinsed with phosphate buffered saline (PBS, pH 7.2), to remove surface blood, which can add difficulty to histological processing.

### Histological quantification of lung metastasis by thin sectioning and hematoxylin and eosin staining

The majority of the lung tissue not utilized for RT-qPCR (4 smaller right lobes; superior, middle, inferior and post-caval, Supplemental Figure 1) were placed into a cassette and immersed in a 10% acetate buffered formalin solution (0.2L 37% formaldehyde, 1.8L distilled H_2_O, and 46.1g Na acetate-3H_2_O) for 24 hours for fixation. The tissue was then rinsed 3 times in 70% ethanol and stored in 70% ethanol until paraffin embedding. For infiltration (embedding), tissues were first dehydrated, cleared with xylene and infiltrated (embedded, Supplemental Table 2). All dehydration, hydration, and xylene steps were conducted in a fumehood for proper air ventilation. Tissues were embedding in paraffin wax using embedding rings, then placed at 4°C for 15 minutes to solidify. Thin sections (5μm) were cut using a fully automated microtome (Leica, RM2255). Cut sections were placed in a 42°C water bath and put on Fisherbrand Superfrost Plus Microscope slides (Fisher Scientific, cat#22-037-246). Slides were dried in a 37°C oven overnight before staining with H&E.

Slides containing paraffin sections were placed in a glass slide holder. Slides were first deparaffinized, then rehydrated in ethanol (Supplemental Table 2). Slides were then deionized in H_2_O before staining (Supplemental Table 3), then dehydrated in ethanol and xylene, in advance of cover slip placement. Before a coverslip was put on the slide, a small drop of Permount (xylene based) was put on the coverslip, the coverslip was angled so that it could gently fall onto the slide, then forceps were used to help guide cover slip in place, allowing the Permount to cover the entire section and to force out air bubbles. Once slides were mounted, they were left to dry overnight in the fumehood.

Images of three sections per block (1-from the first ¼ of block, 1-from the middle of block, and 1-from the last ¼ of block) were captured on the Zeiss Axio Imager Z1 W/ Color and Monochrome camera at 2.5x magnification. To capture the entire lung section, each slide had 4-12 images captured (depending on size of tissue). Images were stitched together using Adobe Photoshop and then imported into the Image J program for metastasis quantification. Each metastatic colony on the lung was Freehand circled and pixelated, then the entire area of the lung was Freehand circled and pixelated, and % metastasis was calculated (Supplemental Figure 1). The total percent metastasis per lung was calculated by averaging percent metastasis of the three quantified thin sections.

### Lung processing for RNA isolation and RNA extraction

The harvested left lung lobe (Supplemental Figure 2) was placed into a petri dish and minced with surgical blades. Other previously described protocols for RNA extraction require additional dissociation and digestions steps (22, 23); however, our protocol includes some modifications, where pure, high-quality RNA can be extracted from lung tissue pieces that are preserved in TRIzol (ThermoFisher, Life Technologies, cat#15596018). Minced pieces were transferred to a microfuge tube and 1mL of TRIzol was added. The tube was vortexed to ensure the minced pieces were dispersed and immersed in TRIzol. At this step, tubes can be flash frozen and stored in a −80°C freeze for later RNA isolation.

Additionally, a naïve mouse lung lobe was spiked with increasing numbers of MDA-MB-231 cells to generate a standard curve for the RT-qPCR-based method. Six tubes containing minced lung tissue and 1mL of TRIzol were prepared and then 0, 10, 100, 1 000, 100 000, or 1 000 000 MDA-MB-231 cells were added to the tubes in 100μL of PBS.

To prevent RNA degradation, it is critical to wear gloves when purifying RNA and handling RNA samples. It is also essential to use RNase/DNase-free certified solutions and plastics and pipette tips need to have filters. Tubes containing TRIzol and tissue were thawed and vortexed until minced tissue was dispersed. In a modification from the manufacturer’s protocol, twice as much chloroform (ThermoFisher) as recommended (400μL) was added to the tubes. The tubes were vortexed for another 10 minutes or until tissue was dissolved. The tubes were centrifuged at 10000xg for 10 minutes for phase separation. The top clear aqueous layer (350μL) was harvested and combined with 350 μl of 70% Ethanol (1:1) and applied to Purelink RNA columns (ThermoFisher, Life Technologies, cat#12183025). RNA was then purified following manufacturer’s protocol and incorporated the addition of DNase (ThermoFisher, Life Technologies, cat#12185010) to eliminate DNA contamination. RNA should be stored at −80°C. RNA concentrations were measured on a Spectramax M2 spectrophotometer (Molecular Devices). Absorbance (A) values at 230, 260 and 280nm were determined and are also an assessment of purity. The RNA samples should have an A260/A280 of ~2.0 and an A260/A230 of 2.0-2.2. Higher A260/230 ratios may indicate organic compound (i.e. TRIzol) contamination and those samples should not be used. RNA samples should also be assessed by gel to confirm that they are not degraded (28). After measuring absorbance of RNA, cDNA should be generated within 7 days to limit changes due to RNA degradation.

### Human-specific GAPDH RT-qPCR

RNA was converted to cDNA using iScript (Bio-Rad, cat#1708890) following the manufacturer’s protocol. Briefly, 0.25μg of RNA of each sample was added to 2μl of 5X iScript reaction mix and RNase free water was added to 10μL of total reaction volume and mixed with a pipette tip. The reactions (in 8-strip PCR tubes) were incubated in a thermal cycler following the manufacturer’s protocol (5 minutes at 25°C, 20 minutes at 46°C, 1 minute at 95°C, hold at 4°C optional). Notably, due to potential low levels of residual DNase contamination, it is important to store cDNA at −20°C, avoid long-term storage beyond a month, and keep cDNA samples on ice when being utilized in RT-qPCR reactions.

Before use in RT-qPCR reactions, the cDNA samples were diluted 1/10 in the 8-strip PCR tubes. Using a multichannel (helps reduce variability between samples due to pipetting error), 4μL was dispensed into 384-well qPCR plate (BioRad). Subsequently, the Ssoadvanced Universal SYBR Green Supermix (BioRad, cat# 172-5270) mastermix containing 5μL of the Ssoadvanced Universal SYBR Green Supermix and 1μL of target primers (from 4μM stock containing forward and reverse primers, was diluted to final working concentration of 0.4μM in the qPCR reaction), according to manufacturer’s protocol. Ten μl of total reaction was loaded per well in duplicate. The sequences for the human-specific and mouse GAPDH primers were generated using the National Center for Biotechnology Information (NCBI) Primer-Basic Local Alignment Search Tool (BLAST, Supplemental Table 1). The RT-qPCR was performed on a CFX384 Touch Real-Time PCR Detection System (Bio-Rad, cat#1855485). Supplemental Table 4 outlines the reaction conditions. Other Real-time PCR detection systems can be utilized.

## DATA AVAILABILITY

All data are within the manuscript and Supplementary Information file.

## FUNDING AND ADDITIONAL INFORMATION

Support was provided by grant funding to PM from the Canadian Institutes of Health Research (CIHR, PJT 162313). MLD is supported by CGS-D award from the CIHR, a Nova Scotia Health Research Foundation studentship, a Nova Scotia graduate scholarship, and a Killam Laureate scholarship. KMC was supported by a CGS-D award from CIHR and by the DeWolfe Graduate Award from the Dalhousie Medical Research Foundation, and a studentship from the Beatrice Hunter Cancer Research Institute and the Canadian Imperial Bank of Commerce.

<u>36948</u>

## ABBREVIATIONS

ALDH1A3: Aldehyde dehydrogenase 1A3
BLAST: Basic Local Alignment Search Tool
BLI: bioluminescence imaging
cDNA: complimentary DNA
FFPE: formalin-fixed paraffin-embedded
GAPDH: glyceraldehyde 3-phosphate dehydrogenase
GFP: green fluorescent protein
H&E: hematoxylin and eosin
HER2: human epidermal growth factor receptor 2
MPRIP: myosin phosphatase rho Interacting protein
MRI: magnetic resonance imaging
NCBI: National Center for Biotechnology Information
NOD/SCID: nonobese diabetic/severe combined immunodeficiency
PTGER2: prostaglandin E receptor-2
RT-qPCR: reverse transcription-quantitative polymerase chain reaction
RNA: ribonucleic acid

